# Function and Cryo-EM structures of broadly potent bispecific antibodies against multiple SARS-CoV-2 Omicron sublineages

**DOI:** 10.1101/2022.08.09.503414

**Authors:** Ping Ren, Yingxia Hu, Lei Peng, Luojia Yang, Kazushi Suzuki, Zhenhao Fang, Meizhu Bai, Liqun Zhou, Yanzhi Feng, Yongji Zou, Yong Xiong, Sidi Chen

## Abstract

The SARS-CoV-2 variant, Omicron (B.1.1.529), rapidly swept the world since its emergence. Compared with previous variants, Omicron has a high number of mutations, especially those in its spike glycoprotein that drastically dampen or abolish the efficacy of currently available vaccines and therapeutic antibodies. Several major sublineages of Omicron evolved, including BA.1, BA.1.1, BA.2, BA.2.12.1, BA.3, BA.4/5, and BA.2.75, which rapidly changing the global and regional landscape of the pandemic. Although vaccines are available, therapeutic antibodies remain critical for infected and especially hospitalized patients. To address this, we have designed and generated a panel of human/humanized therapeutic bispecific antibodies against Omicron and its sub-lineage variants, with activity spectrum against other lineages. Among these, the top clone CoV2-0213 has broadly potent activities against multiple SARS-CoV-2 ancestral and Omicron lineages, including BA.1, BA.1.1, BA.2, BA.2.12.1, BA.3, BA.4/5, and BA.2.75. We have solved the cryo-EM structure of the lead bi-specific antibody CoV-0213 and its major Fab arm MB.02. Three-dimensional structural analysis shows distinct epitope of antibody - spike receptor binding domain (RBD) interactions and reveals that both Fab fragments of CoV2-0213 can simultaneously target one single spike RBD or two adjacent ones in the same spike trimer, further corroborating its mechanism of action. CoV2-0213 represents a unique and potent broad-spectrum SARS-CoV-2 neutralizing bispecific antibody (nbsAb) against the currently circulating major Omicron variants (BA.1, BA.1.1, BA.2, BA.2.12.1, BA.2.75, BA.3, and BA.4/5). CoV2-0213 is primarily human and ready for translational testing as a countermeasure against the ever-evolving pathogen.

---

The SARS-CoV-2 pathogen rapidly disseminates globally, with a constantly evolving landscape in the pandemic ^1^. Numerous variants of concern (VoCs) lineages emerged and continue to evolve (**Fig. S1a**). The Omicron variant (B.1.1.529) rapidly led to multiple new waves globally ^2^. Omicron (sub)variants harbor a high number of mutations, especially in the spike (S) glycoprotein, and clustered in the receptor-binding domain (RBD) (**Fig. S1b-c**). The original Omicron lineage has evolved multiple distinct sub-lineages, such as BA.1, BA.1.1, BA.2, BA.2.12.1, BA.3, BA.4/5, and BA.2.75, which carry various critical mutations (**Fig. S1b-c; Supplemental discussion).** These mutations and (sub)variants drastically decrease the efficacy of currently available vaccines and FDA-approved or emergency-use authorized (EUA) monoclonal antibody-based therapies ^2^. Therefore, it is critical to rapidly develop countermeasures such as new antibodies against these Omicron sublineages. As echoed in recently authorized bi-valent vaccines (Pfizer/BioNTech) and Moderna), it has gain increasing traction to target both Omicron and ancestral lineages that are substantially different, for which a bispecific antibody can achieve with one arm targeting Omicron and the other targeting ancestral lineage.

Among our recently generated Omicron RBD-directed neutralizing monoclonal antibodies (MB.02, MB.08, PC.03) ^3^, all have strong binding activity against Omicron BA.1, while one of these mAbs, MB.02, maintains high activity against BA.2 (0.003 μg/mL, low single-digit ng/mL EC50) (**Fig. S1d**). To enable broad activity against VoCs, we combined Clone13A^4^, a previously generated SARS-CoV-2 RBD-directed neutralizing monoclonal antibody that was potent against SARS-CoV-2 WA-1 and Delta^4^ but had low binding activity against Omicron BA.1 and BA.2 (**Fig. S1e**), with three Omicron mAbs to engineer five bispecific antibodies using an IgG1 knob-into-hole bispecific CrossMab antibody technique ^5^ (**Methods**) (**Fig. 1a**). The resultant bsAb clones were named CoV2-0208, CoV2-0203, CoV2-0803, CoV2-0213, and CoV2-0813. Coomassie-stained SDS-PAGE analysis indicated that all bispecific antibodies were successfully expressed with expected size with high purity after Protein A beads purification (**Fig. 1b, S1f**). Thereafter, antibody titration assays were performed by ELISA and showed strong reactivity of these bsAbs (along with S309, Sotrovimab ^6^) to the RBDs of SARS-CoV-2 WA-1, Delta, Omicron BA.1 and Omicron BA.2 (**Fig. S2a-b**). The remaining bsAbs only showed substantially lower EC_50_(s) to one or two of SARS-CoV-2-RBD(s) (**Fig. S2a**). Meanwhile, our lead bsAbs, CoV2-0213, and CoV2-0813, also exhibited strong competition with angiotensin-converting enzyme 2 (ACE2) for binding to a range of SARS-CoV-2 RBDs (**Fig. S3a**).

**Figure 1.**
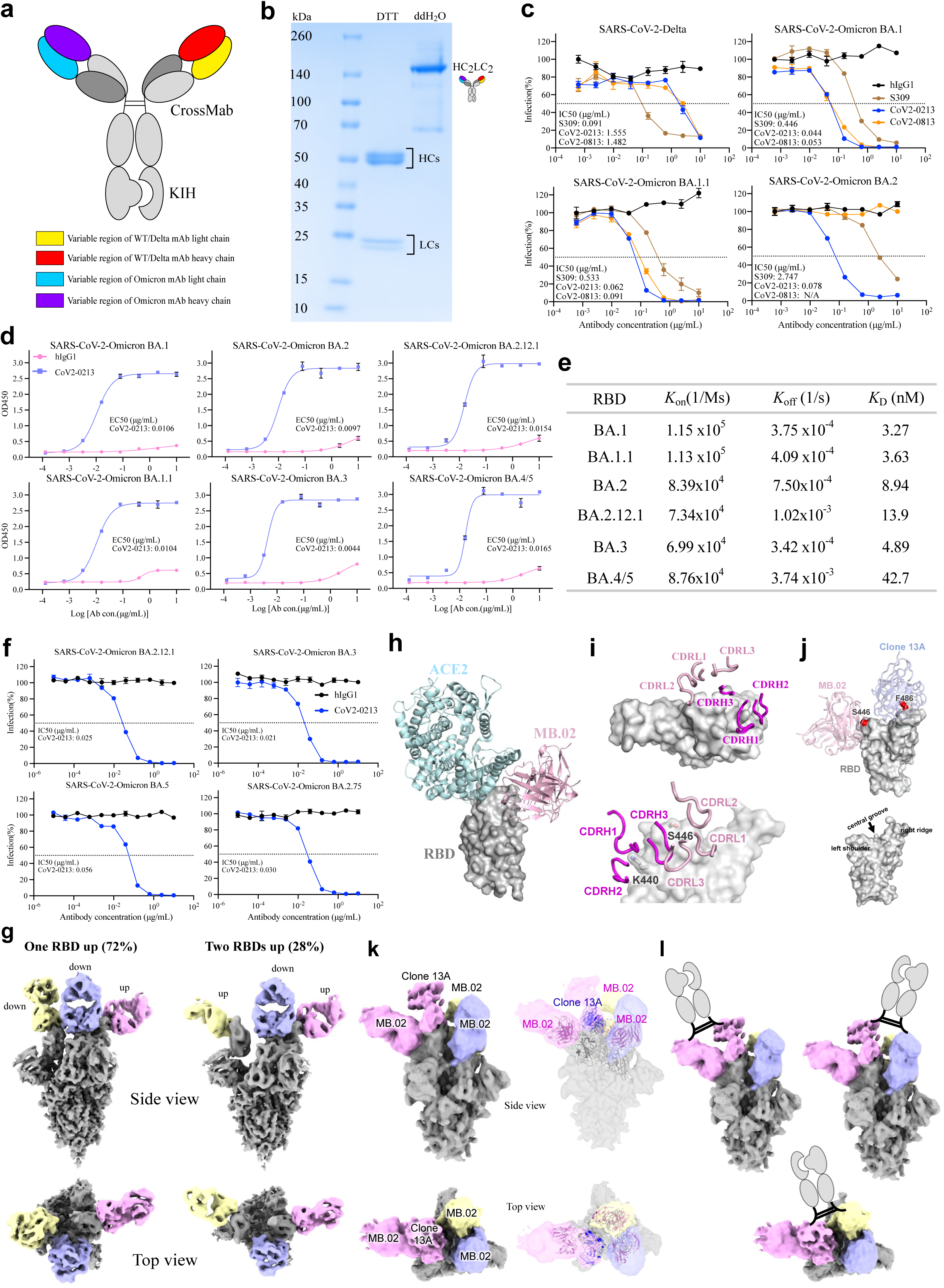
Design, purification, neutralizing activities, and cryo-EM structures of a broadly potent human bispecific antibody CoV2-0213 against circulating Omicron subvariants. **a)** A scheme of bsAb design. Antibody domains are colored according to their architecture. **b)** Coomassie-stained SDS-PAGE analysis of purified bsAb CoV2-0213. The antibody samples are analyzed under reducing condition (+DTT) and nonreducing condition (+ddH_2_O). **c)** Neutralization curves of two lead bsAbs with S309 against pseudotyped virus of SARS-CoV-2 Delta and Omicron sublineages BA.1, BA.1.1, and BA.2. Serial dilutions of bsAb were added to test its neutralizing activity against indicated pseudotyped virus. The IC50 was determined by log (inhibitor) response of nonlinear regression and is displayed as the mean ± s.e.m. **d)** ELISA binding curves of CoV2-0213 with RBD proteins of SARS-CoV-2 Omicron sublineages BA.1, BA.1.1, BA.2, BA.2.12.1, BA.3, and BA.4/5. The EC50 was determined by log (agonist) response of nonlinear regression and is displayed as the mean ± s.e.m.. **e)** The summary statistics of binding affinity of CoV2-0213 to the RBDs of Omicron sublineages as determined by BLI. **f)** Neutralization curves of CoV2-0213 against pseudotyped virus of Omicron sublineages BA.2.12.1, BA.3, BA.5 and BA.2.75. Serial dilutions of bsAb were added to test its neutralizing activity against indicated pseudovirus. The IC50 was determined by log (inhibitor) response of nonlinear regression and is displayed as the mean ± s.e.m.. **g)** Cryo-EM structures of MB.02 Fab fragment in complex with spike trimer in two different conformations, with one RBD up (left) or two RBDs up (right). Fab molecules are shown in different colors, and the spike is shown as dark gray. The corresponding particle distribution of each spike trimer conformation is shown. **h)** Overlay of the structures of human ACE2 (PDB 7wpb) and MB.02 Fab onto the same spike RBD by superimposing the RBD regions. **i)** The binding conformations of the three CDRH and the three CDRL loops of MB.02 Fab on spike RBD. Upper panel, top view; lower panel, side view. The omicron BA.1.1.529 mutation N440K and G446S, which is located at the MB.02 binding interface and inserted between CDRH1 and CDRH2 loops of MB.02 Fab, is shown in stick representation. The nomenclature of the top surface of spike RBD is illustrated in the middle inset. **j)** Overlay of the structures of the humanized clone 2 (Clone 13A) and MB.02 Fab fragments on the same spike RBD. S446 at the MB.02-bound interface of Omicron BA.1/3 spike RBD, and F486 at the Clone 13A-bound interface of WT and Omicron BA.1/2/3 spike RBD, are shown as red spheres. The nomenclature of the top surface of spike RBD is illustrated in the lower panel. **k)** Cryo-EM reconstruction (left panel) and fitted models (right panel) of the CoV2-0213 antibody-bound Omicron BA.5 spike trimer with one RBD up. Fab molecules are shown in different colors, and the spike is shown in dark gray. The up RBD has both MB.02 and Clone 13A Fab fragments bound at distinct interfaces. The other two RBDs in down conformation have the MB.02 Fab bound. The models of the Fab variable domains (MB.02, pink; Clone 13A, blue; RBD, black) are fitted into the 3D reconstruction. **l)** Cartoon illustrations of the possible bivalent binding of one CoV2-0213bsAb onto a single or two adjacent spike RBDs in the same S trimer. The Fc fragment and hinge region of the IgG are shown in the cartoon.

Next, we performed neutralization assays with HIV-1-based pseudoviruses (**Fig. S3b**), a widely used assay by the field that is well-correlated with authentic virus assays ^7–10^. S309 retained neutralization activity against BA.1 and BA.1.1 but its activity against BA.2 dropped 30-fold (**Fig. 1c**). In contrast, CoV2-0213, displayed broad-spectrum neutralization ability to a range of SARS-CoV-2 Omicron VOCs (**Fig. 1c, 1f**). Specifically, CoV2-0213 remained potent in neutralizing seven of Omicron sublineages, BA.1, BA.1.1, BA.2, BA.2.12.1, BA.3, BA.4/5, and BA.2.75 with IC_50_ values of 0.044, 0,062, 0.078, 0.025, 0.021, 0.056, and 0.030 µg/mL, respectively (**Fig. 1c, 1f**). The bsAb CoV2-0213 is an order-of-magnitude more potent than S309/Sotrovimab in neutralization against BA.1 and BA.1.1, and ∼78x more potent than S309/Sotrovimab against BA.2 (**Fig. 1c**). Meanwhile, two other bispecific antibodies also exhibit potent Omicron-specific neutralization (**Fig. 1c**). CoV2-0203 showed high potency in neutralizing the three of Omicron sublineages, although it showed relatively weak neutralization ability (1.5-fold weaker) in comparison to CoV2-0213 (**Fig. S4c**). CoV2-0208 showed high potency against BA.1 and BA.1.1, but its activity against BA.2 is on par with S309 (**Fig. S4c**). Both CoV2-0208 and CoV2-0203 lost neutralization activity against the Delta variant. CoV2-0213 has strong neutralization potency against Omicron BA.1, BA.1.1, and BA.2 and maintains reasonable activity against Delta.

To evaluate antibody cross-reactivity, we further evaluated eight human coronaviruses RBD proteins, which included six Omicron sublineages (BA.1, BA.1.1, BA.2, BA.2.12.1, BA.3, and BA.4/5) and two β-coronaviruses (SARS-CoV and MERS-CoV). The results showed that CoV2-0213 has a broad and strong binding activity to all assayed Omicron RBDs (**Fig. 1d**), with moderate binding to SARS-CoV RBD (**Fig. S2c**). Biolayer interferometry (BLI) results revealed that CoV2-0213 displays high single-molecule affinity, at single-digit nanomolar Kd against Omicron BA.1 (Kd= 3.27nM) and BA.1.1 (Kd= 3.63nM) (**Fig. 1e, S5a**). Meanwhile, CoV2-0213 retains high affinity with low nanomolar Kd against Omicron BA.2 (Kd = 8.94 nM),_BA.2.12.1 (Kd = 13.9 nM), BA.3 (Kd = 4.89 nM) and BA.4/5 (Kd = 42.7 nM), respectively (**Fig. 1e, S5b**).

We determined the cryo-EM structures of MB.02 Fab in complex with the ectodomain of Omicron BA.1 spike trimer (S trimer) at ∼3.2 Å resolution (**MB.02::BA.1 Complex; Table S1**). Two major S trimer conformation states were detected, one with one RBD up (72%) and the other with two RBDs up (28%) (**Fig. 1g**). In both conformations, the S trimer is bound with three Fab molecules, one per RBD. The binding of MB.02 Fab makes the spike RBD more flexible, especially in up conformation (**Fig. S6a**). MB.02 Fab mainly contacts a flexible loop region at the left shoulder region of spike RBD. The MB.02 Fab-binding interface on spike RBD has no overlap with the ACE2-binding interface, although there could be slight steric clashes between the bound ACE2 and MB.02 (**Fig. 1h**). All six complementarity determining regions (CDRs) of the MB.02 Fab participate in RBD interactions (**Fig. 1i**). The MB.02-binding interface contains two mutations (N440K and G446S), with N440K contacting CDRH1 and CDRH2 of MB.02, and G446S interacting with the CDRL2 loop (**Fig. 1i**, lower panel). We analyzed the RBD mutations in BA.2.12.1, BA.3, and BA.4/5 that are directly located at the CoV2-0213 binding interface. The mutation G446S which could be important for MB.02 binding (**Fig. 1j**), is present in BA.3 but not in BA.2.12.1 and BA.4/5, while F486V which may disrupt the Clone 13A interaction, is present in BA.4/5 only (**Fig. 1j**). These differences may explain the enhanced binding affinity of CoV2-0213 bsAb to different Omicron subvariants, although other spike RBD mutations may also have indirect allosteric effects on the RBD conformation at the CoV2-0213 binding interface.

To confirm structural result and further evaluate the possible antibody evasion caused by RBD mutations, we additional performed neutralization assays with pseudotyped site-specific alanine mutant (triple-alanine Omicron BA.1 mutant). The results demonstrated artificial K440A, S446A, V486A Omicron mutant (**Fig. S4a**) can result in strong resistance to CoV2-0213, indicating above three residues are the key epitopes of CoV2-0213, consistent with our structural findings.

Taken together, CoV2-0213 exhibited significant enhanced activities to a wide range of assayed Omicron subvariants compared to its parental monoclonal antibodies^3^, which prompts us to investigate its distinct mechanism of action. One Fab arm of CoV2-0213, Clone 13A, the humanized clone 2 antibody for WT spike ^4^, mainly interacts with the right ridge of spike RBD and would not lead to any steric clash with a bound MB.02, suggesting they could target the same RBD simultaneously (**Fig. 1j**).

To further investigate how CoV2-0213 bsAb binds to the spike antigen, we determined the cryo-EM structure of CoV2-0213 bsAb in complex with the recently emerged and prevalent BA.5 Omicron subvariant at a resolution of 7.7Å (unmasked and unsharpened) (**CoV-0213::BA.5 Complex; Table S1**). Among the cryo-EM particles we collected, only one spike conformation with one RBD up was detected (**Fig. 1k**). From that, we further identified a subset (∼24%) of particles with density for two Fab fragments on the same RBD in the up conformation, and one Fab fragment each for the other two RBDs in the down conformation (**Fig. 1k, left panel**). The density for the Fab-bound up-RBD fitted well with the overlaid model of spike RBD with the MB.02 and Clone 13A Fab fragments^4^, while the density of the Fab-bound down RBDs matched with the model of the MB.02 Fab-bound RBD (**Fig. 1k, right panel**). This suggests three MB.02 and one Clone 13A Fabs bound to the same S trimer, potentially from three 0213 bsAbs with one having both Fab arms bound. Considering the flexible nature of the hinge region of an IgG, it is possible that the two Fab arms of a CoV2-0213 bsAb can target one single spike RBD or two adjacent ones in the same trimer simultaneously (**Fig. 1l**). Alternative interpretation is that four Fabs from four different CoV2-0213 bsAbs bound to spike trimer. Regardless of modes of binding, the cryo-EM data revealed epitope co-engagement mechanism and supported the enhanced engagement of the Omicron spike and functional activity of CoV2-0213.

In summary, we have generated five distinct SARS-CoV-2-targeting human nbsAbs, and the lead nbsAb, CoV2-0213, exhibited potent and broad-spectrum activities ability to a range of SARS-CoV-2 VoCs, including various Omicron sublineages and ancestral lineages. Cryo-EM structure demonstrated that the two Fabs arms of CoV2-0213 can fully engage the same RBD simultaneously. CoV2-0213 is primarily human and ready for translational testing.

## Acknowledgments

We thank various members from our labs for discussions and support. We thank staffs from various Yale core facilities for technical support. We thank all staffs at the Laboratory for BioMolecular Structure (LBMS) at Brookhaven National Lab for support in cryo-EM data collection. The LBMS is supported by the DOE Office of Biological and Environmental Research (KP1607011). We thank Drs. Klein, L Chen, Müschen and others for providing equipment and related support. We thank various support from Department of Genetics; Institutes of Systems Biology and Cancer Biology; Dean’s Office of Yale School of Medicine and the Office of Vice Provost for Research.

## Funding

This work is supported by DoD PRMRP IIAR (W81XWH-21-1-0019) and discretionary funds to SC and NIH R01 AI163395 to YX.

## Institutional Approval

All recombinant DNA (rDNA) and biosafety work were performed under the guidelines of Yale Environment, Health and Safety (EHS) Committee with approved protocols (Chen 15-45, 18-45, 20-18, 20-26).

## Author Contributions

PR: design of the bispecific antibodies, construct generation, cloning, antibody expression, ELISA, data analysis, figure prep, writing

YH: Cryo-EM, structure studies, writing

LP: construct generation, cloning, neutralization, data analysis, figure prep

LY, KS, MB, YF, ZZ: assisting experiments and data collection

ZF, LZ: construct generation, cloning, resources

YX: Cryo-EM, structure studies, funding, supervision, writing

SC: conceptualization, overall design, funding, supervision, writing

## Supplemental discussion

The SARS-CoV-2 pathogen rapidly disseminates globally, with a constantly evolving landscape in the pandemic ^1^. To date, the COVID-19 pandemic has infected nearly 600 million confirmed individuals and caused over 6.4 million deaths (https://www.worldometers.info/coronavirus/). One of the most concerning features of the virus is that the pathogen SARS-CoV-2 continues to evolve. Numerous mutant lineages emerged, some became dominant while some diminished, leading to multiple waves across the world. The Omicron variant (B.1.1.529), initially reported in South Africa in late 2021, rapidly became the predominant variant circulating in many countries and led the fourth pandemic wave globally ^2^. As compared with the genome sequences of ancestral variants of concerns (VOCs), the Omicron lineage has harbored a high number of genomic mutations, especially in the spike (S) glycoprotein and clustered in the receptor-binding domain (RBD). These mutations drastically decrease the efficacy of currently available vaccines and FDA-approved or emergency-use authorized (EUA) monoclonal antibody-based therapies ^2–8^.

As of today, the original Omicron lineage has evolved into multiple distinct sub-lineages, such as BA.1, BA.2, BA.3, and BA.4/BA.5 according to WHO weekly epidemiological update^9–11^. BA.1.1 (a subvariant of BA.1) has an additional R346K mutation and caused large regional outbreaks in Canada with BA.1 during the first quarter of 2022^12^. BA.2.12.1 (a subvariant of BA.2), which carried two additional alterations (L452 and S704) on top of BA.2, has emerged and once become the dominant variant in the US and certain other regions ^13^. In addition, BA.4/BA.5, which shared an identical sequence of the spike protein, contained additional mutations (Del69-70, L452R, F486V, and R493Q) compared to BA.2, and become dominant in several major regions of the world, including the US and China^14^. These phenomena indicate rapid dynamics of the viral evolution, leading to substantial changes in infectivity, antigenic escape, reduction of vaccine efficacy, and increased possibility of repeat infections^5, 6, 13, 15, 16^. Although vaccines are available, therapeutic antibodies remain critical for infected and especially hospitalized patients. However, few clinical monoclonal antibodies can retain substantial activities against all Omicron sublineages ^13^, urging for more therapeutic options^14, 17^. Therefore, it is critical to rapidly develop countermeasures such as new antibodies against these sublineages.

In this correspondence study, we, therefore, set out to develop new candidates of monoclonal therapeutic antibodies that can counter multiple Omicron sub-lineages. To avoid potential immune escape and infection by SARS-CoV-2 mutational variants, cocktail strategies (i.e., administration of two human monoclonal antibodies) were previously designed and reported, which had been demonstrated that could largely increase efficacy and maximally prevent viral escape ^18–20^. However, the cocktail strategy requires the production of multiple molecules. On the contrary, the bispecific antibody (bsAb) strategy can simultaneously target two different antigens or antigenic sites according to its structural design. In addition, bsAb can benefit from the synergistic effects of the two binder Fab arms ^21^. An effective dual-binder bsAb can also save the manufacturing process cost by half as compared to a cocktail of two mono-specific mAbs, which can be substantial in translational and clinical stage development. Furthermore, the dual-targeting concept has already been successfully studied as an effective strategy in the treatment of cancer and inflammatory disorders ^22–24^. In addition, several studies have illustrated that bispecific antibodies could potently enhance breadth and potency than parental monoclonal antibodies ^25–27^. Due to extensive mutations in the spike protein, Omicron sublineages are drastically different from the variants in the ancestral lineages (e.g., wildtype, WT, Wuhan-1, WA-1) and earlier variants (such as Alpha variant, Beta variant, Delta variant).

We have generated five distinct SARS-CoV-2-targeting nbsAbs (CoV2-0208, CoV2-0203, CoV2-0803, CoV2-0213, and CoV2-0813), and evaluated their functional properties against the ancestral and Omicron (sub)lineages. We included the clinically relevant mAb S309 (Sotrovimab ^28^) in the experiment, which has been in human use under Food and Drug Administration (FDA) issued emergency use authorization (EUA) in the US, and similarly in Europe, UK, Japan, and Australia. The mAb S309 retained neutralization activity against both Omicron BA.1 and BA.1.1 (BA.1+R346 mutation) with half-maximum inhibitory concentration (IC_50_) values of 0.44 and 0.53 µg/mL, respectively. However, its neutralization activity against Omicron BA.2 dropped 30-fold to an IC_50_ value of 2.75µg/mL. Of note, the EUA of S309/Sotrovimab was recently withdrawn by FDA in March 2022 due to the high prevalence of the BA.2 variant in many states in the US.

To characterize our bispecific antibody candidates for their reactivity against multiple SARS-CoV-2 lineages, we performed a series of assays, including ELISA, Biolayer interferometry (BLI), and neutralization. To better understand the mechanism of action of the bsAb CoV-0213, we performed cryo-electron microscopy (cryo-EM) studies to solve the three-dimensional (3D) structures of the bsAb and its major Fab arm (MB.02) in complexes with the Omicron spike variants (we previously reported the cryo-EM structure of the other Fab arm, 13A in complex with WT spike^29^). We first sought to map the molecular basis for the broad specificity of MB.02 and its interaction with the RBD epitope and determined the cryo-EM structures of MB.02 Fab in complex with the ectodomain of Omicron BA.1 spike trimer (S trimer) at ∼3.2 Å resolution. To further investigate how CoV2-0213 bsAb binds to the spike antigen, we determined the cryo-EM structure of CoV2-0213 bsAb in complex with the recently emerged and prevalent BA.5 Omicron subvariant. The omicron BA.1 spike RBD maintains an overall architecture similar to that of the WT spike from the ancestorial lineage, with minor local conformational changes primarily at the mutation sites.

The lead bispecific antibody, CoV2-0213, exhibited significantly or even enhanced binding affinity and neutralizing efficacy to a wide range of assayed Omicron subvariants compared to its parental monoclonal antibodies^30^, suggesting its promise as a potent and more cost-effective candidate as compared to using two single mAbs. To further investigate how CoV2-0213 bsAb binds to the spike antigen, we determined the cryo-EM structure of CoV2-0213 bsAb in complex with the recently emerged and prevalent BA.5 Omicron subvariant (CoV-0213::BA.5 Complex). The flexible Fc region could not be visualized in the cryo-EM reconstruction and therefore we could not identify which bound MB.02 Fab and Clone 13A Fab were from the same CoV2-0213 bsAb. This structure showed that three MB.02 and one Clone 13A Fabs bound to the same S trimer, potentially from three 0213 bsAbs with one having both Fab arms bound. Considering the flexible nature of the hinge region of an IgG, it is possible that the two Fab arms of a CoV2-0213 bsAb can target one single spike RBD or two adjacent ones in the same trimer simultaneously. It is also possible the observed four bound-Fab fragments are from four different CoV2-0213 bsAbs. Regardless of which mode the bsAb binds to the spike RBD, our observation explains the enhanced engagement of the Omicron spike and neutralization effect by CoV2-0213.

Our lead nbsAb, CoV2-0213, exhibited potent and broad-spectrum activities ability to all major Omicron sublineages (BA.1, BA.1.1, BA.2, BA.2.12.1, BA.3 BA.4/BA.5, and BA.2.75). While the Omicron sublineages continue to evolve, the recent branches such as XBB, BQ.1.1, BU.1, BR.2, CA.2, and CJ.1 all stem from B.1.1.529. In fact, the dominant variant BQ.1.1 is a sublineage of BA.5, and XBB is a sublineage of BA.2. Therefore, CoV2-0213 has good coverage against these Omicron sublineages. While CoV2-0213 it is less potent against the ancestral lineages such as WT/WA1, Alpha and Delta, those variants had been long disappeared since late 2021 - early 2022 in the natural course of viral evolution. CoV2-0213 has good coverage against Omicron sublineages. Our cryo-EM structure demonstrated that the two Fabs arms of CoV2-0213 can fully engage the same SARS-CoV-2 spiker trimer simultaneously. Furthermore, Omicron BA.1 triple-alanine mutant abolished neutralization to CoV-0213, indicating K440, S446, and V486 are the key residues that retained the neutralizing activities of CoV2-0213. In summary, CoV2-0213 is primarily human and ready for translational testing as a countermeasure against the ever-evolving pathogen.

## Methods

### Cloning and expression of SARS-CoV2-specific human bispecific antibodies

Bispecific antibodies were cloned and expressed using methods similar to those previously described ^29^, utilizing the Fab regions from four of our previously developed fully human or largely humanized monoclonal antibodies, Clones MB.02, MB.08, PC.03^30^ and Clone 13A^29^. In brief, indicated mAb variable regions for each bispecific antibody were amplified and subcloned into separate mammalian expression vectors using Gibson assembly^29^. To express recombinant bispecific antibodies, four expression vectors were transiently transfected into Expi293 cells using ExpiFectamine 293 transfection kit according to the manufacturer’s protocol (ThermoFisher). Antibody containing cell culture supernatants were harvested after 5 days of cultivation in shake flasks, then secreted bispecific antibodies were collected and purified by affinity chromatography using rProtein A Sepharose Fast Flow beads (Cytiva, Cat: #17127901). Purified bispecific antibodies were inspected using SDS-PAGE and stored at −80°C after further usage.

### SARS-CoV-2 pseudotyped virus generation and neutralization assay

The SARS-CoV-2 pseudotyped virus was produced as previously described ^30^. Briefly, pseudotyped virus containing cell culture supernatant was harvested after 2 days of co-transfection of HEK-293T cells with a spike-expressing plasmid and env-deficient HIV-1 backbone vectors, then clarified by centrifugation and stored at −80°C after further usage. To determine the neutralizing activity of the bispecific antibody, serially diluted antibodies were incubated with pseudotyped virus at 37°C for 1 hour, then co-cultured with HEK-293T-hACE2 cells for overnight. Finally, the signal was evaluated after 48 hours by detection of GFP expression in the HEK-293T-hACE2 cells using Attune NxT Acoustic Focusing Cytometer (ThermoFisher) or BD Symphony Flow Cytometry.

### Antibody binding quantification and ACE2 competition assay

The binding of bispecific antibodies were quantified by ELISA and ACE2 competition assays, as previously described ^30^. The recombinant MERS-CoV RBD (Cat. No. 50-201-9463), SARS-CoV RBD (Cat. No. 50-196-4017), SARS-CoV-2 RBD wild type (WA-1) (Cat. No. 40592-V08B), SARS-CoV-2 Delta RBD (Cat. No.40592-V08H90), SARS-CoV-2 Omicron BA.1 RBD (Cat. No.40592-V08H121), SARS-CoV-2 Omicron BA.1.1 RBD (Cat. No. SPD-C522j-100ug), SARS-CoV-2 Omicron BA.2 RBD (Cat. No.SPD-C522G-100ug), SARS-CoV-2 Omicron BA.2.12.1 RBD (Cat. No. SPD-C522Q-100ug), SARS-CoV-2 Omicron BA.3 RBD (Cat. No. SPD-C522I-100ug), SARS-CoV-2 Omicron BA.4/5 RBD (Cat. No. SPD-C522R-100ug) used in ELISA quantification were purchased from Thermo, Sino Biological and AcroBiosystems, respectively^30^. The ACE2 (Cat. No.10108-H08H) used in the ACE2 competition assay was purchased from Sino Biological ^29^.

### Bispecific antibody binding affinity measurement

The binding affinity of antibodies with RBD was performed previously by Octet RED96e (ForteBio) using bio-layer interferometry (BLI). 25ng/μl of purified bispecific antibody was immobilized onto AHC biosensors (ForteBio) with recombinant Omicron RBDs acting as the analyte with serial dilutions. Kd values were calculated using Data Analysis HT 10 (ForteBio) with a 1:1 Langmuir binding model.

### Cryo-EM sample preparation and data collection

The omicron B.1.1.529 spike trimer (Sino Biological Cat: 40589-V08H26) at a final concentration of 0.6 mg/mL was mixed with MB.02 Fab at a molar ratio of 1:1.5 on ice for 30mins. The omicron BA.5 spike trimer (ACROBiosystems SPN-C522e-50ug) at a final concentration of 0.6 mg/mL was mixed with the bispecific antibody 0213 at a molar ratio of 1:1 on ice for 30mins. Then 3 μl of the mixture was applied to a Quantifoil-Au-2/1-3C grid (Quantifoil) pretreated by glow-discharging at 20 mA for 1 min. The grid was blotted at 18 °C with 100% humidity and plunge-frozen in liquid ethane using FEI Vitrobot Mark IV (Thermo Fisher). The grids were stored in liquid nitrogen until data collection.

Images were acquired on an FEI Titan Krios electron microscope (Thermo Fisher) equipped with a Gatan K3 Summit direct detector in super-resolution mode, at a calibrated magnification of 81,000× with the physical pixel size corresponding to 1.07 Å. Detailed data collection statistics for the Fab-spike trimer complexes are shown in a supplemental table. Automated data collection was performed using SerialEM ^31^.

### Cryo-EM data processing

A total of 5373 movie series were collected for the complex of MB.02 Fab with omicron BA.1 S trimer. A total of 2408 movie series were collected for the complex of CoV2-0213 bsAb with the omicron BA.5 S trimer. Motion correction of the micrographs was carried out using RELION ^32^ and contrast transfer function (CTF) estimation was calculated using CTFFIND4 ^33^. Particles were picked automatically by crYOLO ^34^, followed by 2D and 3D classifications without imposing symmetry. The 3D classes with different S trimer conformations were then processed separately by consensus 3D refinement and CTF refinement. For each state of the complex, local masked 3D classification without image alignment was performed focusing on one Fab-RBD region, and the best class of particles was selected for consensus refinement of the whole complex. For the complex of CoV2-0213 bsAb with the omicron BA.5 S trimer, repeated consensus 3D refinement and subsequent local masked 3D classification without image alignment, focusing on the up RBD with the two associated Fabs, were performed to identify the class of S trimers with both MB.02 and Clone 13A Fabs bound to the same up RBD. Subsequently, multibody refinement was performed as described above for the rigid body containing the focused region. The 3D reconstruction of the other Fab-RBD regions were obtained with the same procedure. The final resolution of each reconstruction was determined based on the Fourier shell correlation (FSC) cutoff at 0.143 between the two half maps ^35^. The final map of each body was corrected for K3 detector modulation and sharpened by a negative B-factor estimated by RELION ^36^, and then merged in Chimera for deposition. The local resolution estimation of each cryo-EM map is calculated by RELION^32^(**Fig. S6b and S6c**).

### Model building and refinement

The structure of SARS-CoV-2 omicron BA.1 spike trimer (PDB 7qo7) was used as an initial model and docked into the spike trimer portion of the cryo-EM maps using Chimera ^37^. The initial model of MB.02 Fab was generated by homology modeling using SWISS-MODEL ^38^, and then docked into the Fab portions of the cryo-EM maps using Chimera ^37^. The initial models were subsequently manually rebuilt in COOT ^39^, followed by real-space refinement in PHENIX ^40^. The final models with good geometry and fit to the map were validated using the comprehensive cryo-EM validation tool implemented in PHENIX ^41^. All structural figures were generated using PyMol (http://www.pymol.org/) and ChimeraX ^42^.

### Schematic illustrations

Schematic illustrations were created with Affinity Designer.

## Reporting summaries

### Statistics

For all statistical analyses, confirmed that the items mentioned in the NPG reporting summary are present in the figure legend, table legend, main text, or Methods section.

### Standard statistical analysis

All statistical methods are described in figure legends and/or supplementary Excel tables. **Source data and statistics** were provided in a supplemental excel table.

### Software and code

#### Data collection

ELISA data were recorded by a microplate reader (Perkin Elmer) (no version number). Antibody binding kinetics for anti-spike mAbs were evaluated by BLI on an Octet RED96e instrument (FortéBio) at room temperature (version 12). Flow cytometry data were collected by ThermoFisher Attune and/or BD FACS Aria flow cytometers.

Cryo-EM data were acquired on a FEI Titan Krios electron microscope (Thermo Fisher) equipped with a Gatan K3 Summit direct detector in super-resolution mode, at a calibrated magnification of 81,000× with the physical pixel size corresponding to 1.070 Å. Detailed data collection statistics for the Fab-spike trimer complexes are shown in a supplemental table. Automated data collection was performed using SerialEM (v3.8).

#### Data analysis

Data analysis was performed using the following software/code:

Flow data were analyzed by FlowJo v.10.7

Standard biological assays’ data were analyzed in Prism (v8 or v9).

BLI data was analyzed by using Octet Analysis Studio Software 10.0.

Motion correction of the micrographs was carried out using RELION (v4.0) and contrast transfer function (CTF) estimation was calculated using CTFFIND4 (v4.1).

Particles were picked automatically by crYOLO (v1.8.1), followed by 2D and 3D classifications without imposing symmetry. The 3D classes with different S trimer conformations were then processed separately by consensus 3D refinement and CTF refinement.

The final map of each body was corrected for K3 detector modulation and sharpened by a negative B-factor estimated by RELION and then merged in Chimera for deposition.

The structure of the ectodomain of SARS-CoV-2 Omicron BA.1 spike trimer (PDB 7qo7) was used as an initial model and docked into the spike trimer portion of the cryo-EM maps using Chimera (v1.15).

The initial structural model of MB.02 Fab was generated by homology modeling using SWISS-MODEL (online https://swissmodel.expasy.org), and then docked into the Fab portions of the cryo-EM maps using Chimera.

The initial models were subsequently manually rebuilt in COOT (0.9.7), followed by real-space refinement in PHENIX (v1.19).

The final models with good geometry and fit to the map were validated using the comprehensive cryo-EM validation tool implemented in PHENIX. All structural figures were generated using PyMol (v1.3) (online http://www.pymol.org/) and ChimeraX (v1.2).

#### Data and resource availability

All data generated or analyzed during this study are included in this article and its supplementary information files. Specifically, source data and statistics for regular experiments are provided in an excel file of **Source data and statistics**. The models of the mAb:Spike complexes have been deposited in the wwPDB with pending accession codes (1uSpike-MB.02, 8DZH; 2uSpike-MB.02, 8DZI). The cryo-EM maps of the mAb:Spike complexes have been deposited in EMDB with pending accession codes (1uSpike-MB.02, EMD-27798; 2uSpike-MB.02, EMD-27799; 1uSpike-CoV2-0213, EMD-27800). Other materials and data are available either through public repositories or via reasonable requests to the corresponding authors to the academic community.

#### Code availability

No custom code was used in this study.

### Life sciences study design

#### Sample size determination

The sample size was determined according to the lab’s prior work or similar approaches in the field. For most cases, at least biological triplicate experiments (n >= 3) were performed unless otherwise noted. Details on sample size for experiments were indicated in methods and figure legends.

#### Data exclusions

No data were excluded.

#### Replication

All experiments were done with replicates. Experimental replications were indicated in detail in the methods section and each figure panel’s legend. All key experiments were successfully replicated in the lab to ensure the rigor of the findings.

#### Randomization

Each sample was randomly allocated into experimental or control groups.

#### Blinding

The experiments were not blinded.

### Reporting for specific materials, systems and methods

#### Antibodies used

Custom Antibodies were generated in this study, where dilutions were often serial titrations (i.e. a number of dilutions as specified in each figure and/or legend.)

COV2-0208 (generated in this study)

COV2-0203 (generated in this study)

COV2-0803 (generated in this study)

COV2-0213 (generated in this study)

COV2-0813 (generated in this study)

S309 (Sotrovimab)

MB.02

MB.08

PC.03

Clone13A

Commercial antibodies used for staining were the following, with typical dilutions listed:

Mouse anti-Human lgGl Fc Secondary Antibody, HRP Thermo Fisher Cat#A-10648, 1:2000

InVivoMAb human lgGl isotype control BioXcell Cat#BE0297.

Goat Anti-Mouse lgG H&L (HRP) Abcam ab6789, Abcam Cat#ab6789, 1:5000.

#### Antibody validation

Commercial antibodies were validated by the vendors and re-validated in-house as appropriate through antigen-specific experiments. Custom antibodies were validated by specific antibody-antigen interaction assays, such as ELISA. Isotype controls were used for antibody validations.

Commercial antibody info and validation info where applicable:

https://www.thermofisher.com/antibody/product/Mouse-anti-Human-IgG1-Fc-Secondary-Antibody-clone-HP6069-Monoclonal/A-10648

https://bxcell.com/product/invivomab-human-igg1-isotype-control/

https://www.abcam.com/goat-mouse-igg-hl-hrp-ab6789.html

### Eukaryotic cell lines

#### Cell line source(s)

All commercial cell lines were originally acquired from commercial vendors (ATCC, ThermoFisher). HEK293T (Thermo Fisher), and 293FT-hACE2 (gifted from Dr. Bieniasz’ lab) cell lines were used in this study.

#### Authentication

Cell lines were authenticated by the original commercial vendors.

#### Mycoplasma contamination

All the cell lines used here tested negative for mycoplasma contamination.

#### Commonly misidentified lines (See ICLAC register)

No commonly misidentified line was used in the study.

### Flow Cytometry

#### Plots

Confirm the checkboxes of requirements.

### Methodology

#### Sample preparation

Sample prep description in the Methods section.

#### Instrument

Flow cytometric analysis was performed on a BD FACSAria II or ThermoFisher Attune™ NxT.

#### Software

FlowJo v.10.7.1 was used for flow cytometry data analysis.

#### Cell population abundance

N/A

#### Gating strategy

Cells were gated by FSC/SSC plot. To distinguish between positive and negative boundaries of the stained cells, negative control samples were analyzed and utilized as background.

#### Gating example figure

Confirm that a figure exemplifying the gating strategy is provided in the Supplementary Information.

## Editorial Policy Items

### Additional policy considerations

#### Macromolecular structural data

##### Validation report

We have provided an official validation report from wwPDB for all macromolecular structures studied.

##### Biological materials

Other materials and data are available either through public repositories or via reasonable requests to the corresponding authors to the academic community.

---

**Table S1.**
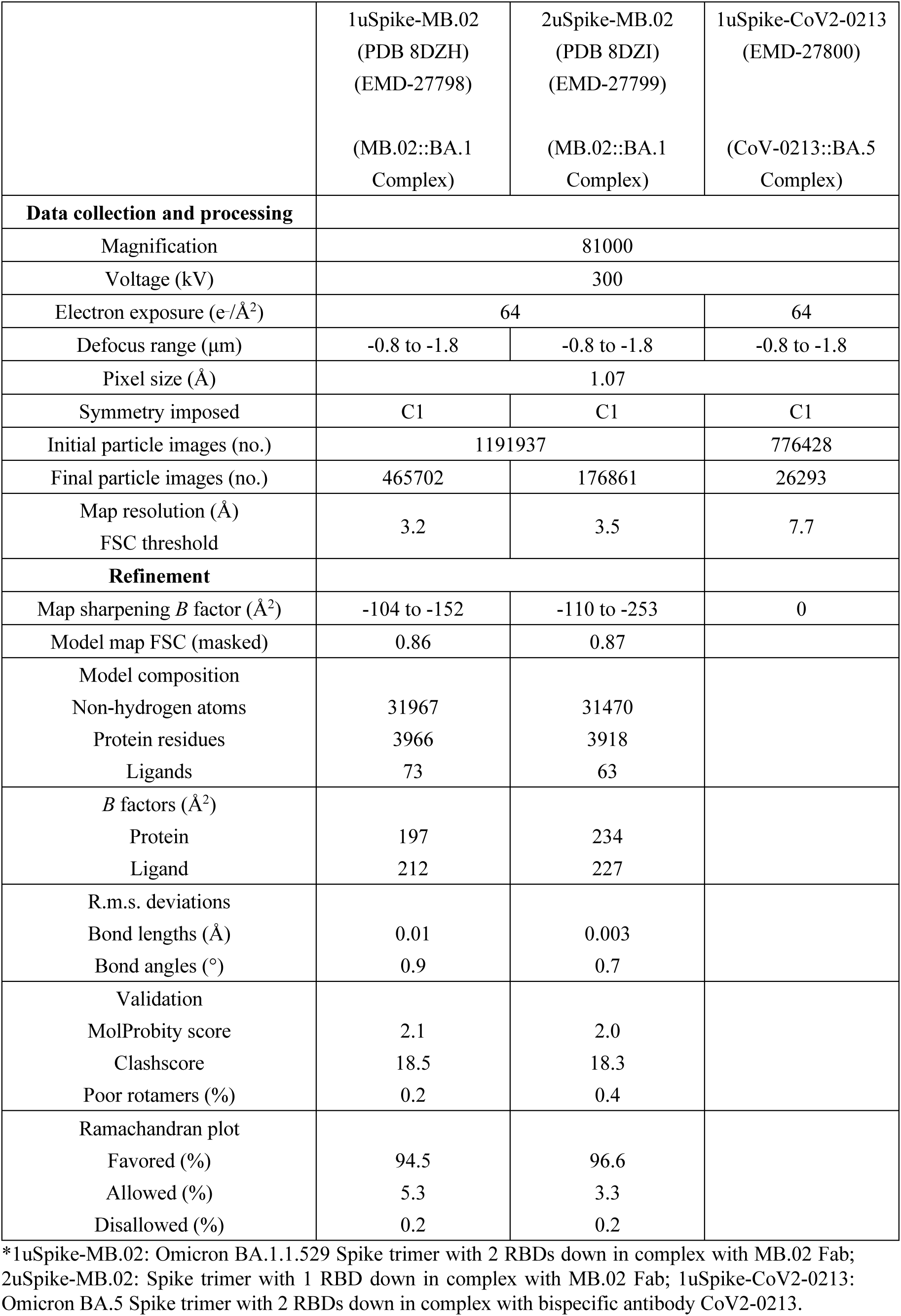
Cryo-EM Data Collection and Refinement Statistics

|  | 1uSpike-MB.02<br>(PDB 8DZH)<br>(EMD-27798)<br>(MB.02::BA.1<br>Complex) | 2uSpike-MB.02<br>(PDB 8DZI)<br>(EMD-27799)<br>(MB.02::BA.1<br>Complex) | 1uSpike-CoV2-0213<br>(EMD-27800)<br>(CoV-0213::BA.5<br>Complex) |
| --- | --- | --- | --- |
| **Data collection and processing** |  |  |  |
| Magnification | 81000 |  |  |
| Voltage (kV) | 300 |  |  |
| Electron exposure (e-/Å<sup>2</sup>) | 64 |  | 64 |
| Defocus range (μm) | -0.8 to -1.8 | -0.8 to -1.8 | -0.8 to -1.8 |
| Pixel size (Å) | 1.07 |  |  |
| Symmetry imposed | C1 | C1 | C1 |
| Initial particle images (no.) | 1191937 |  | 776428 |
| Final particle images (no.) | 465702 | 176861 | 26293 |
| Map resolution (Å)<br>FSC threshold | 3.2 | 3.5 | 7.7 |
| **Refinement** |  |  |  |
| Map sharpening *B* factor (Å<sup>2</sup>) | -104 to -152 | -110 to -253 | 0 |
| Model map FSC (masked) | 0.86 | 0.87 |  |
| Model composition |  |  |  |
| Non-hydrogen atoms | 31967 | 31470 |  |
| Protein residues | 3966 | 3918 |  |
| Ligands | 73 | 63 |  |
| *B* factors (Å<sup>2</sup>) |  |  |  |
| Protein | 197 | 234 |  |
| Ligand | 212 | 227 |  |
| R.m.s. deviations |  |  |  |
| Bond lengths (Å) | 0.01 | 0.003 |  |
| Bond angles (°) | 0.9 | 0.7 |  |
| Validation |  |  |  |
| MolProbity score | 2.1 | 2.0 |  |
| Clashscore | 18.5 | 18.3 |  |
| Poor rotamers (%) | 0.2 | 0.4 |  |
| Ramachandran plot |  |  |  |
| Favored (%) | 94.5 | 96.6 |  |
| Allowed (%) | 5.3 | 3.3 |  |
| Disallowed (%) | 0.2 | 0.2 |  |

\*1uSpike-MB.02: Omicron BA.1.1.529 Spike trimer with 2 RBDs down in complex with MB.02 Fab; 2uSpike-MB.02: Spike trimer with 1 RBD down in complex with MB.02 Fab; 1uSpike-CoV2-0213: Omicron BA.5 Spike trimer with 2 RBDs down in complex with bispecific antibody CoV2-0213.

**Figure S1.**
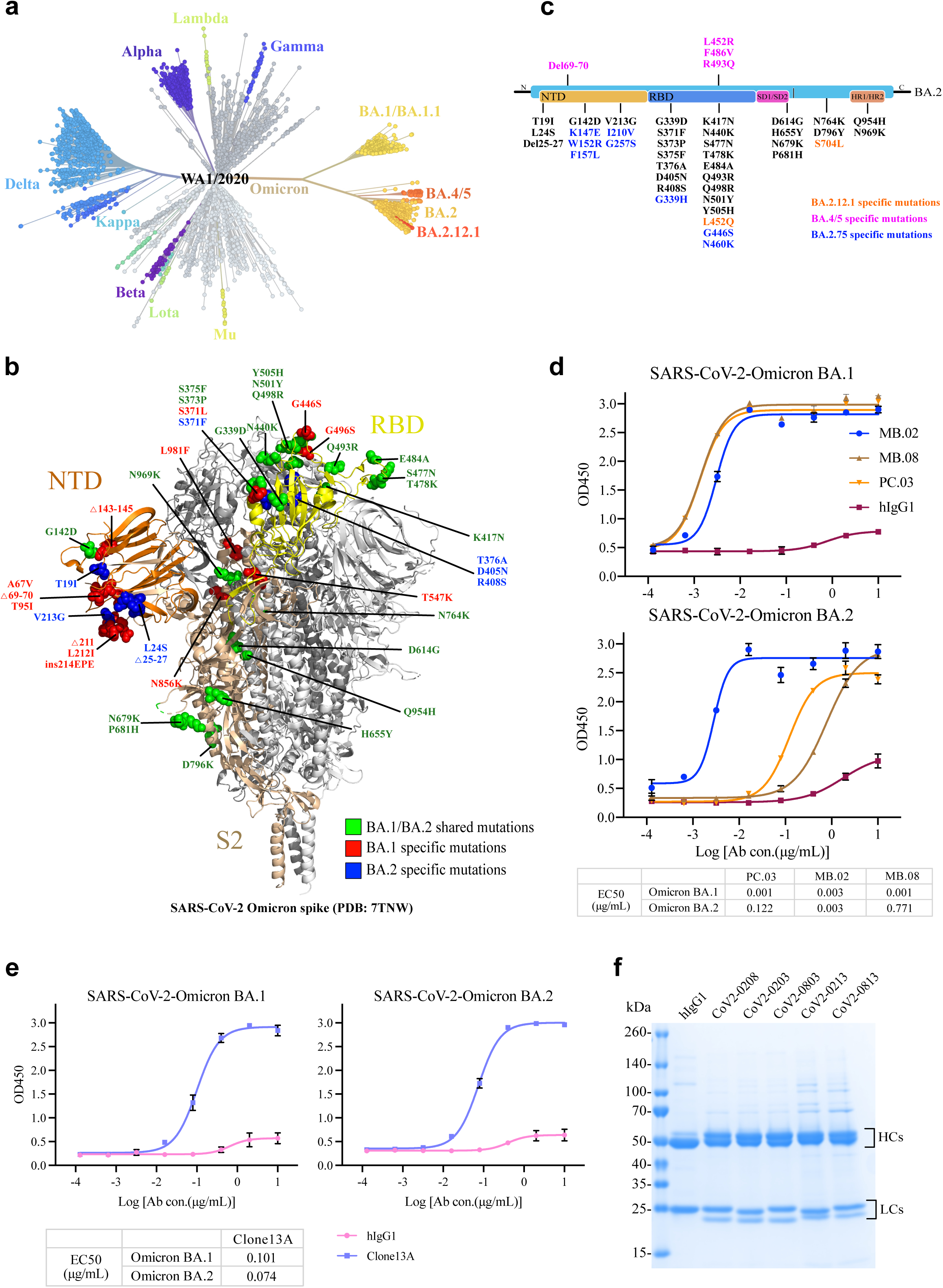
Phylogeny, spike amino acid changes, and antibody evasion of the Omicron subvariant. **(a)** Unrooted phylogenetic tree of Omicron and its subvariants along with other major SARS-CoV-2 variants, figure modified from the Nextstrain (https://nextstrain.org/ncov/gisaid/global/all-time), using data available from the GISAID initiative. **(b)** The structure of the closed prefusion conformation of the Omicron S trimer (PDB: 7TNW) is shown in the ribbon diagram with one protomer colored as NTD in orange, RBD in yellow, and the N-terminal segment of S2 in wheat. All mutations in the Omicron subvariant BA.1 and BA.2 are highlighted in the sphere model. **(c)** Specific spike mutations found in Omicron BA.2.12.1, Omicron BA.2.75 and Omicron BA4/5 are colored compared with the sequence of Omicron BA.2 sublineage (Orange, Omicron BA.2.12.1 specific mutations; Blue, Omicron BA.2.75 specific mutations; Violet, Omicron BA4/BA.5 specific mutations)**. (d, e)** ELISA binding curves of three parental Omicron BA.1 RBD specific mAbs **(d)** and one parental SARS-CoV-2 RBD specific mAb **(e)** to Omicron sublineages RBD domains. The EC50 was determined by log (agonist) response of nonlinear regression and is displayed as the mean ± s.e.m. **(f)** Coomassie-stained SDS-PAGE analysis of five purified resultant bsAbs. The antibody samples are analyzed under reducing condition (+DTT).

**Figure S2.**
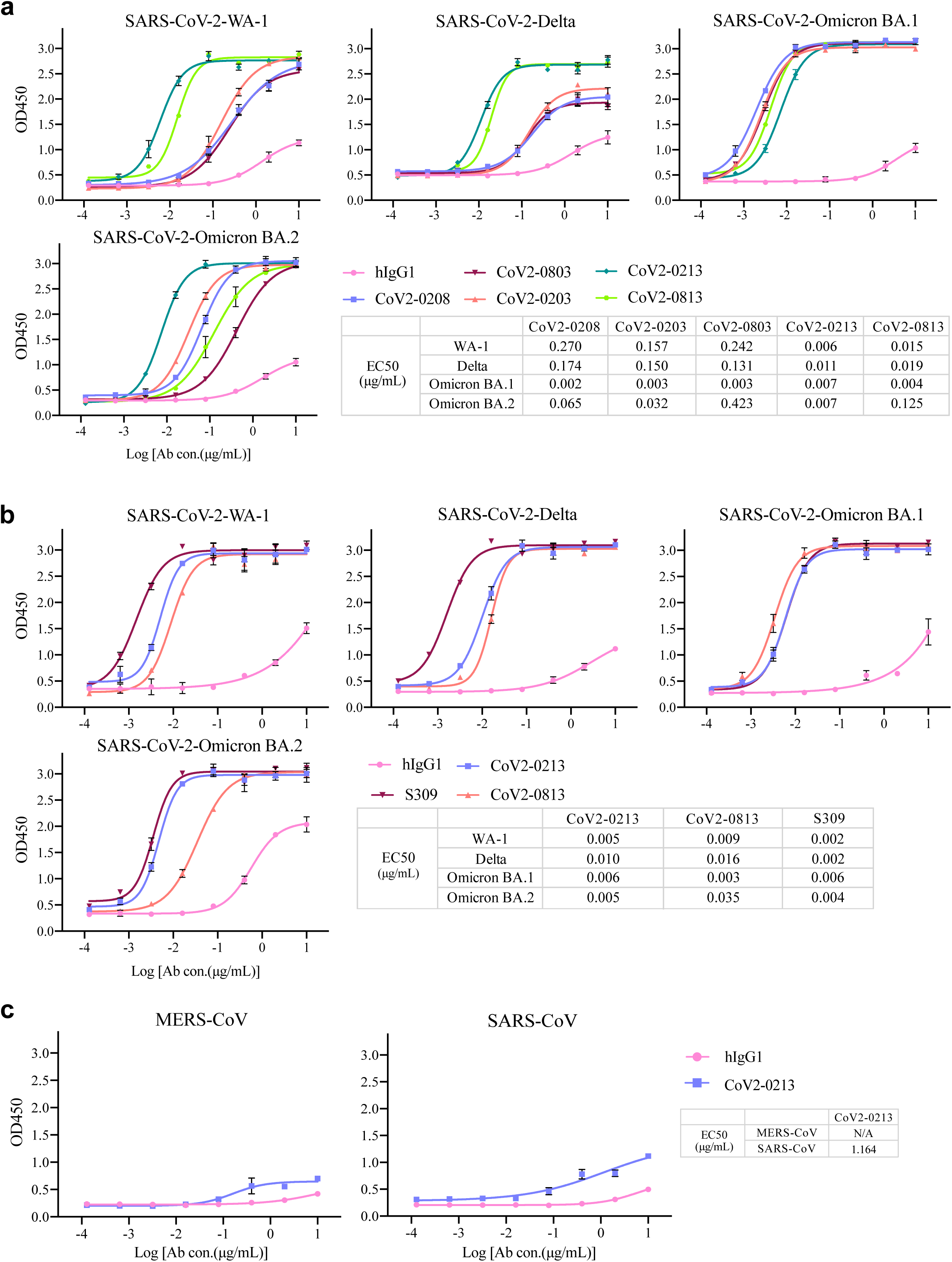
ELISA-based binding activities of resultant bsAbs are superior to those of its parental mAbs. **(a)** ELISA binding curves of five resultant bsAbs with RBD proteins of SARS-CoV-2 WA-1, Delta, Omicron BA.1 and BA.2. The EC50 was determined by log (agonist) response of nonlinear regression and is displayed as the mean±s.e.m. **(b)** ELISA binding curves of two lead bsAbs and S309 with RBD proteins of SARS-CoV-2 WA-1, Delta, Omicron BA.1 and BA.2. The EC50 was determined by log (agonist) response of nonlinear regression and is displayed as the mean±s.e.m. **(c)** ELISA binding curves of CoV2-0213 with RBD proteins of MERS-CoV and SARS-CoV. The EC50 was determined by log (agonist) response of nonlinear regression and is displayed as the mean ± s.e.m.

**Figure S3.**
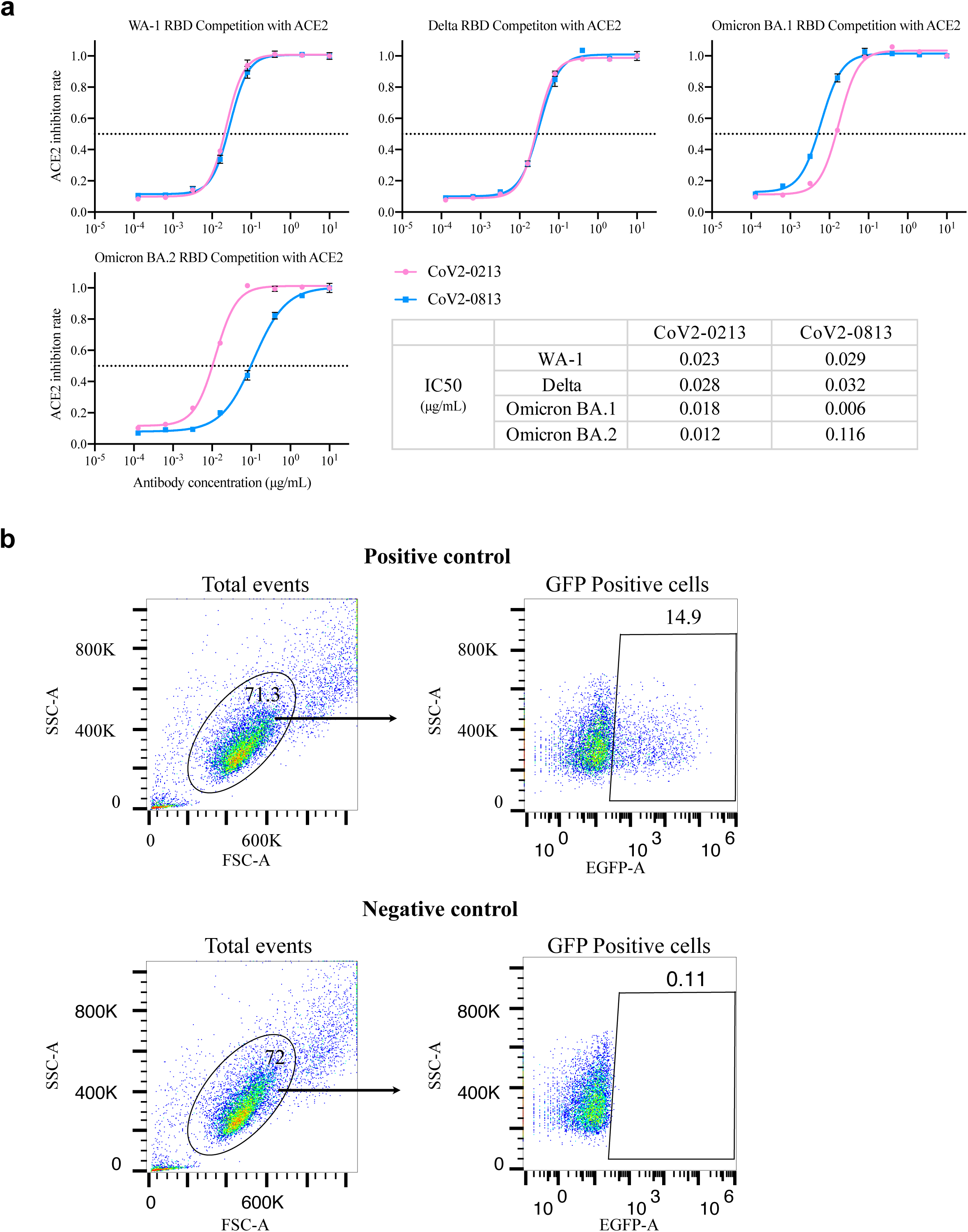
ACE2/RBD binding inhibition for two lead bsAbs and flow gating strategy of neutralization assay. **(a)** Competition ELISA of lead bispecific Abs with human ACE2 for epitope identification to SARS-CoV-2 WT, Delta, Omicron BA.1 and BA.2 RBD proteins. The data were obtained from a representative experiment with three replicates. Data are represented as mean ± s.e.m. IC50 values were calculated from ACE2 competition ELISA assays by fitting a log (inhibitor) response of a nonlinear regression model. **(b)** Representative flow gating plots.

**Figure S4.**
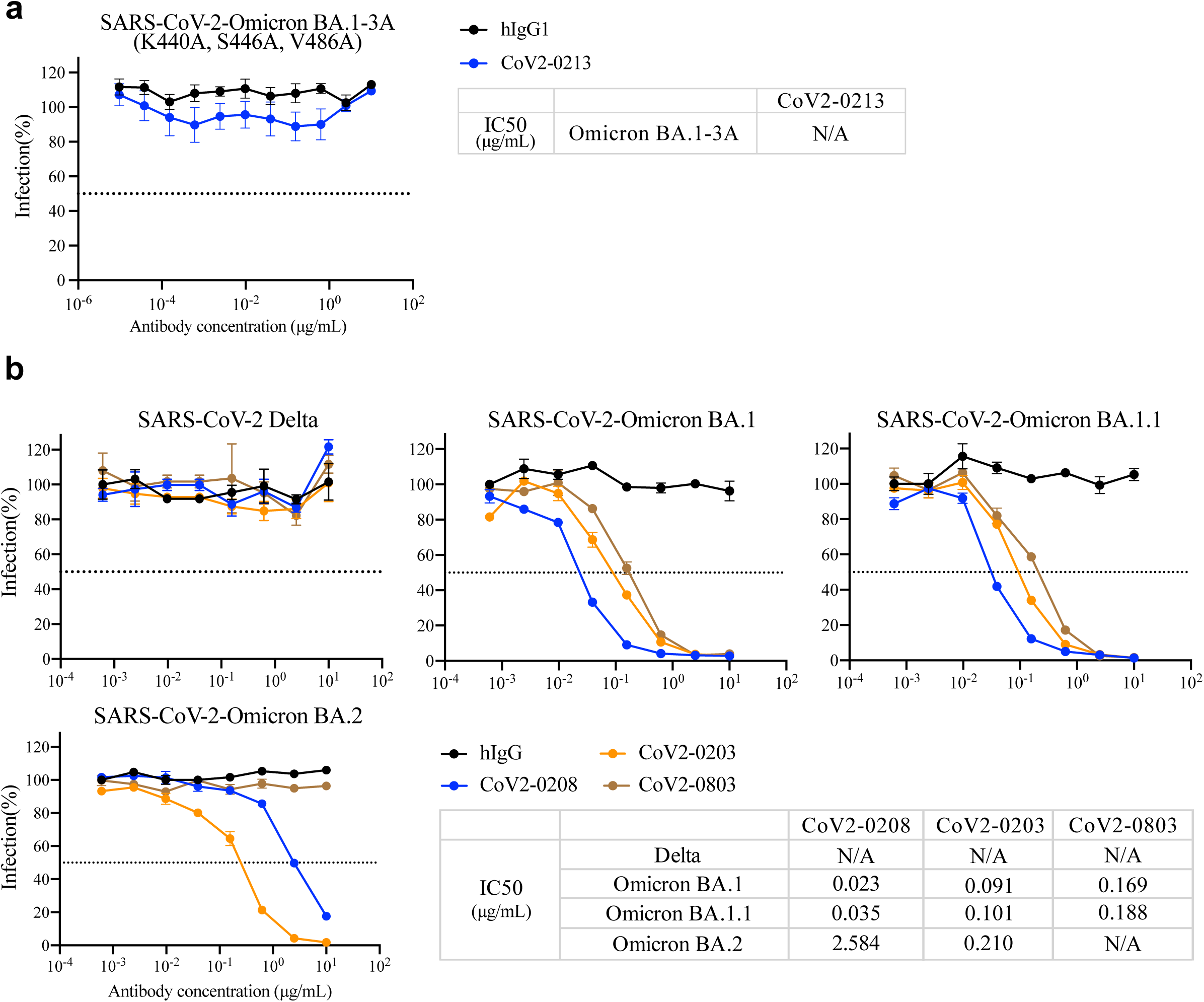
Resistance of SARS-CoV and SARS-CoV-2 subvariants to neutralization by resultant bsAbs. **(a)** Modeling of K440A, S446A and V486A on Omicron BA.1 RBD abolish CoV2-0213 neutralization. Serial dilutions of bsAb were added to test its neutralizing activity against indicated pseudotyped virus. The IC50 was determined by log (inhibitor) response of nonlinear regression and is displayed as the mean ± s.e.m. **(b)** Neutralization curves of remaining bsAbs against pseudotyped virus of ancestorial SARS-CoV-2 subvariants. Serial dilutions of bsAb were added to test its neutralizing activity against indicated pseudotyped virus. The IC50 was determined by log (inhibitor) response of nonlinear regression and is displayed as the mean ± s.e.m.

**Figure S5.**
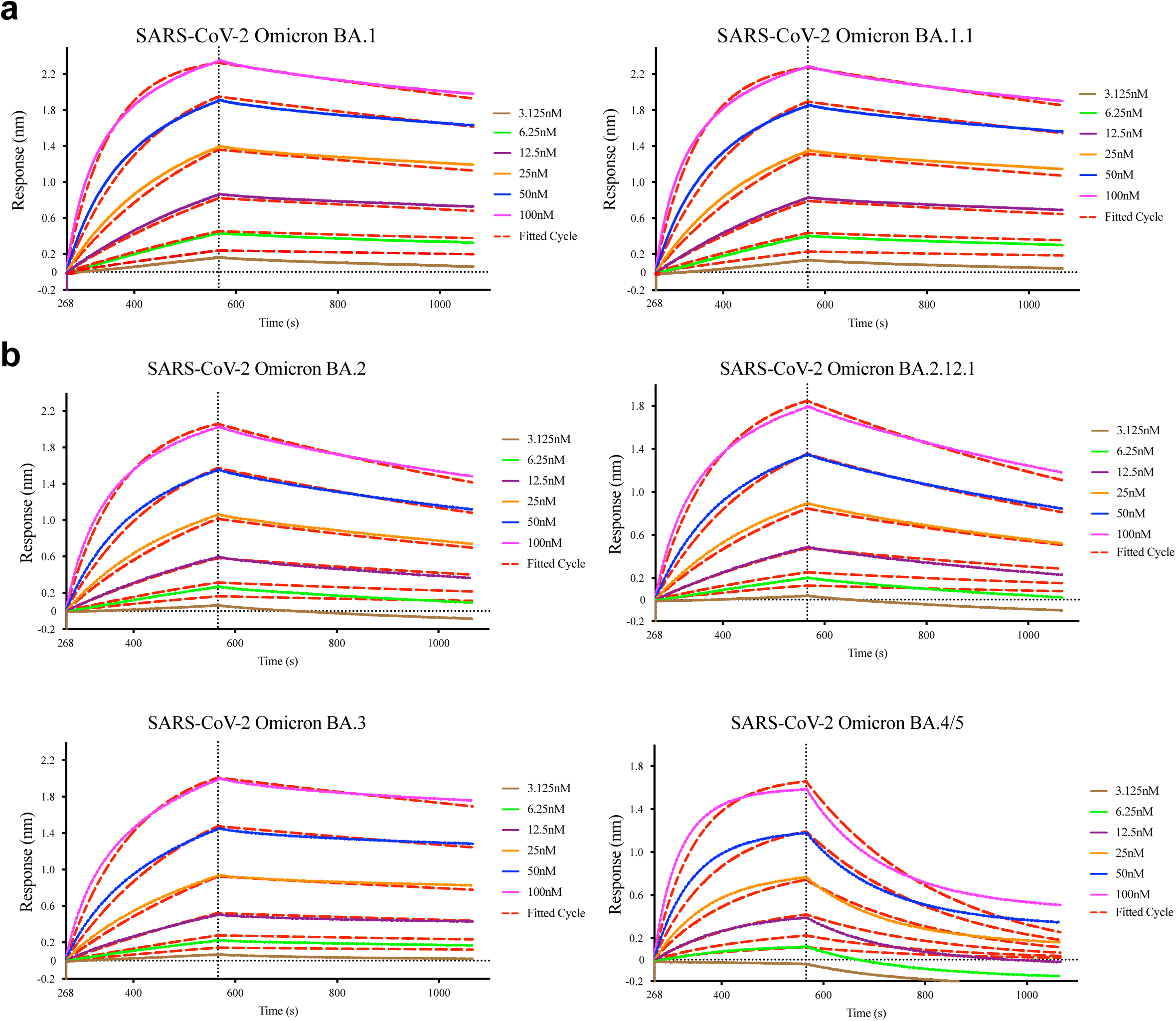
Affinity of the RBDs of SARS-CoV-2 Omicron subvariants to CoV2-0213. **(a)** Binding affinities of the RBDs of Omicron subvariant BA.1 and BA.1.1 to CoV2-0213 as measured by BLI. bsAbs were immobilized on an AHC sensor and tested for real-time association and dissociation of the RBD (BA.1, left panel, BA.1.1, right panel). Global fit curves are shown as red dotted lines. The vertical dashed lines indicate the transition between the association and dissociation phase. **(b)** Binding affinities of the RBDs of Omicron subvariant BA.2, BA.2.12.1, BA.3, and BA.4/5 to CoV2-0213 as measured by BLI. bsAbs were immobilized on an AHC sensor and tested for real-time association and dissociation of the RBD (BA.2, left upper panel, BA.2.12.1, right upper panel, BA.3, left lower panel, BA.4/5, right lower panel). Global fit curves are shown as red dotted lines. The vertical dashed lines indicate the transition between the association and dissociation phase.

**Figure S6.**
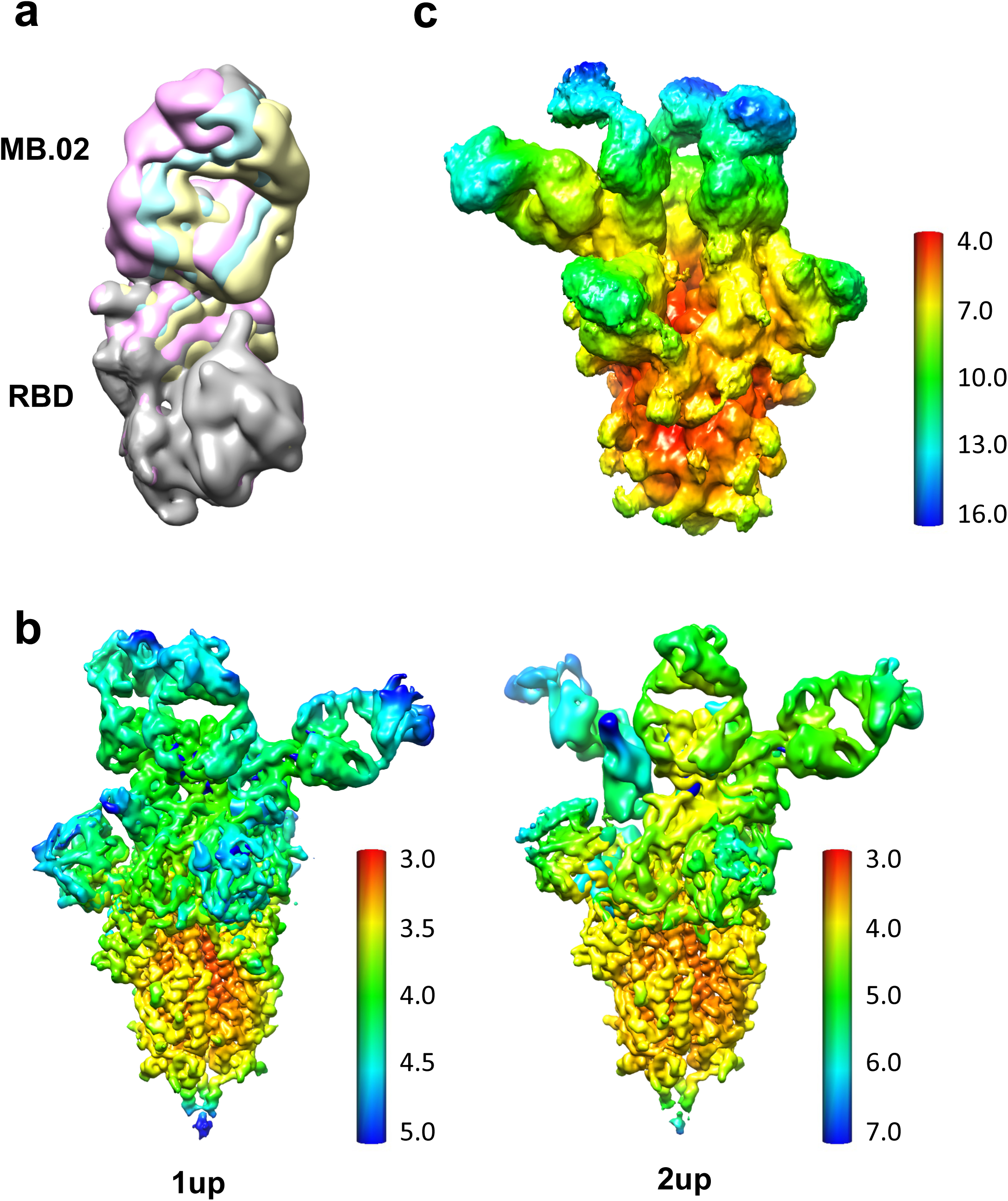
Local flexibility and resolution estimation of the cryo-EM reconstructions. **(a)** Example of one RBD-MB.02 Fab protomer that has flexible conformations. **(b)** Local resolution estimation of the reconstructions of MB.02 Fab in complex with Omicron BA.1 spike trimer of two different conformations. **(c)** Local resolution estimation of the reconstruction of bispecific antibody CoV2-0213 in complex with Omicron BA.5 spike trimer.

## Source data and statistics

Source data and statistics provided in an excel file

## Notes

### Competing Interest Statement

The authors have declared no competing interest.

### Summary of Updates

Added additional data.

## References

1 Acuti Martellucci, C., et al. SARS-CoV-2 pandemic: An overview. Adv Biol Regul 77, 100736 (2020). https://doi.org:10.1016/j.jbior.2020.100736

2 Liu, L. et al. Striking antibody evasion manifested by the Omicron variant of SARS-CoV-2. Nature 602, 676–681 (2022). https://doi.org:10.1038/s41586-021-04388-0

3 P. Ren, L. P., Z. Fang, K. Suzuki, P. Renauer, Q. Lin, M. Bai, L. Yang, T. Li, P. Clark, D. Klein, S. Chen. Potent and specific human monoclonal antibodies against SARS-CoV-2 Omicron variant by rapid mRNA immunization of humanized mice. bioRxiv (2022).

4 Peng, L. et al. Monospecific and bispecific monoclonal SARS-CoV-2 neutralizing antibodies that maintain potency against B.1.617. Nat Commun 13, 1638 (2022). https://doi.org:10.1038/s41467-022-29288-3

5 Schaefer, W. et al. Immunoglobulin domain crossover as a generic approach for the production of bispecific IgG antibodies. Proc Natl Acad Sci U S A 108, 11187–11192 (2011). https://doi.org:10.1073/pnas.1019002108

6 Gupta, A. et al. Early Treatment for Covid-19 with SARS-CoV-2 Neutralizing Antibody Sotrovimab. N Engl J Med 385, 1941–1950 (2021). https://doi.org:10.1056/NEJMoa2107934

7 Bewley, K. R. et al. Quantification of SARS-CoV-2 neutralizing antibody by wild-type plaque reduction neutralization, microneutralization and pseudotyped virus neutralization assays. Nat Protoc 16, 3114–3140 (2021). https://doi.org:10.1038/s41596-021-00536-y

8 Garcia-Beltran, W. F. et al. mRNA-based COVID-19 vaccine boosters induce neutralizing immunity against SARS-CoV-2 Omicron variant. Cell 185, 457–466 e454 (2022). https://doi.org:10.1016/j.cell.2021.12.033

9 Liu, L. et al. Potent neutralizing antibodies against multiple epitopes on SARS-CoV-2 spike. Nature 584, 450–456 (2020). https://doi.org:10.1038/s41586-020-2571-7

10 Nie, J. et al. Quantification of SARS-CoV-2 neutralizing antibody by a pseudotyped virus-based assay. Nat Protoc 15, 3699–3715 (2020). https://doi.org:10.1038/s41596-020-0394-5

## References

3 Cao, Y. et al. Omicron escapes the majority of existing SARS-CoV-2 neutralizing antibodies. Nature 602, 657–663 (2022). https://doi.org:10.1038/s41586-021-04385-3

4 Cele, S. et al. Omicron extensively but incompletely escapes Pfizer BNT162b2 neutralization. Nature 602, 654–656 (2022). https://doi.org:10.1038/s41586-021-04387-1

5 Iketani, S. et al. Antibody evasion properties of SARS-CoV-2 Omicron sublineages. Nature (2022). https://doi.org:10.1038/s41586-022-04594-4

6 Planas, D. et al. Considerable escape of SARS-CoV-2 Omicron to antibody neutralization. Nature 602, 671–675 (2022). https://doi.org:10.1038/s41586-021-04389-z

7 Rossler, A., Riepler, L., Bante, D., von Laer, D. & Kimpel, J. SARS-CoV-2 Omicron Variant Neutralization in Serum from Vaccinated and Convalescent Persons. N Engl J Med 386, 698–700 (2022). https://doi.org:10.1056/NEJMc2119236

8 Zhou, T. et al. Structural basis for potent antibody neutralization of SARS-CoV-2 variants including B.1.1.529. Science, eabn8897 (2022). https://doi.org:10.1126/science.abn8897

9 Evans, J. P. et al. Neutralization of the SARS-CoV-2 Deltacron and BA.3 Variants. N Engl J Med 386, 2340–2342 (2022). https://doi.org:10.1056/NEJMc2205019

10 Hachmann, N. P. et al. Neutralization Escape by SARS-CoV-2 Omicron Subvariants BA.2.12.1, BA.4, and BA.5. N Engl J Med 387, 86–88 (2022). https://doi.org:10.1056/NEJMc2206576

11 Yu, J. et al. Comparable Neutralization of the SARS-CoV-2 Omicron BA.1 and BA.2 Variants. medRxiv (2022). https://doi.org:10.1101/2022.02.06.22270533

12 Brown, P. E., et al. Omicron BA.1/1.1 SARS-CoV-2 Infection among Vaccinated Canadian Adults. N Engl J Med 386, 2337–2339 (2022). https://doi.org:10.1056/NEJMc2202879

13 Wang, Q. et al. Antibody evasion by SARS-CoV-2 Omicron subvariants BA.2.12.1, BA.4, & BA.5. Nature (2022). https://doi.org:10.1038/s41586-022-05053-w

14 Takashita, E. et al. Efficacy of Antibodies and Antiviral Drugs against Omicron BA.2.12.1, BA.4, and BA.5 Subvariants. N Engl J Med (2022). https://doi.org:10.1056/NEJMc2207519

15 Tuekprakhon, A. et al. Antibody escape of SARS-CoV-2 Omicron BA.4 and BA.5 from vaccine and BA.1 serum. Cell 185, 2422–2433 e2413 (2022). https://doi.org:10.1016/j.cell.2022.06.005

16 Cao, Y., et al. BA.2.12.1, BA.4 and BA.5 escape antibodies elicited by Omicron infection. Nature (2022). https://doi.org:10.1038/s41586-022-04980-y

17 Vangeel, L., et al. Remdesivir, Molnupiravir and Nirmatrelvir remain active against SARS-CoV-2 Omicron and other variants of concern. Antiviral Res 198, 105252 (2022). https://doi.org:10.1016/j.antiviral.2022.105252

18 Baum, A. et al. REGN-COV2 antibodies prevent and treat SARS-CoV-2 infection in rhesus macaques and hamsters. Science 370, 1110–1115 (2020). https://doi.org:10.1126/science.abe2402

19 Baum, A. et al. Antibody cocktail to SARS-CoV-2 spike protein prevents rapid mutational escape seen with individual antibodies. Science 369, 1014–1018 (2020). https://doi.org:10.1126/science.abd0831

20 Wang, N. et al. Structure-based development of human antibody cocktails against SARS-CoV-2. Cell Res 31, 101–103 (2021). https://doi.org:10.1038/s41422-020-00446-w

21 Li, C. et al. Broad neutralization of SARS-CoV-2 variants by an inhalable bispecific single-domain antibody. Cell 185, 1389–1401 e1318 (2022). https://doi.org:10.1016/j.cell.2022.03.009

22 Demanet, C., Brissinck, J., De Jonge, J. & Thielemans, K. Bispecific antibody-mediated immunotherapy of the BCL1 lymphoma: increased efficacy with multiple injections and CD28-induced costimulation. Blood 87, 4390–4398 (1996).

23 Li, J. F., Niu, Y. Y., Xing, Y. L. & Liu, F. A novel bispecific c-MET/CTLA-4 antibody targetting lung cancer stem cell-like cells with therapeutic potential in human non-small-cell lung cancer. Biosci Rep 39 (2019). https://doi.org:10.1042/BSR20171278

24 Xiong, M. et al. A Novel CD3/BCMA Bispecific T-cell Redirecting Antibody for the Treatment of Multiple Myeloma. J Immunother 45, 78–88 (2022). https://doi.org:10.1097/CJI.0000000000000401

25 Bournazos, S., Gazumyan, A., Seaman, M. S., Nussenzweig, M. C. & Ravetch, J. V. Bispecific Anti-HIV-1 Antibodies with Enhanced Breadth and Potency. Cell 165, 1609–1620 (2016). https://doi.org:10.1016/j.cell.2016.04.050

26 Frei, J. C. et al. Bispecific Antibody Affords Complete Post-Exposure Protection of Mice from Both Ebola (Zaire) and Sudan Viruses. Sci Rep 6, 19193 (2016). https://doi.org:10.1038/srep19193

27 Moshoette, T., Ali, S. A., Papathanasopoulos, M. A. & Killick, M. A. Engineering and characterising a novel, highly potent bispecific antibody iMab-CAP256 that targets HIV-1. Retrovirology 16, 31 (2019). https://doi.org:10.1186/s12977-019-0493-y

28 Gupta, A. et al. Early Treatment for Covid-19 with SARS-CoV-2 Neutralizing Antibody Sotrovimab. N Engl J Med 385, 1941–1950 (2021). https://doi.org:10.1056/NEJMoa2107934

29 Peng, L. et al. Monospecific and bispecific monoclonal SARS-CoV-2 neutralizing antibodies that maintain potency against B.1.617. Nat Commun 13, 1638 (2022). https://doi.org:10.1038/s41467-022-29288-3

30 P. Ren, L. P., Z. Fang, K. Suzuki, P. Renauer, Q. Lin, M. Bai, L. Yang, T. Li, P. Clark, D. Klein, S. Chen. Potent and specific human monoclonal antibodies against SARS-CoV-2 Omicron variant by rapid mRNA immunization of humanized mice. bioRxiv (2022).

31 Mastronarde, D. N. Automated electron microscope tomography using robust prediction of specimen movements. J Struct Biol 152, 36–51 (2005). https://doi.org:10.1016/j.jsb.2005.07.007

32 Kimanius, D., Dong, L., Sharov, G., Nakane, T. & Scheres, S. H. W. New tools for automated cryo-EM single-particle analysis in RELION-4.0. Biochem J 478, 4169–4185 (2021). https://doi.org:10.1042/BCJ20210708

33 Rohou, A. & Grigorieff, N. CTFFIND4: Fast and accurate defocus estimation from electron micrographs. J Struct Biol 192, 216–221 (2015). https://doi.org:10.1016/j.jsb.2015.08.008

34 Wagner, T. et al. SPHIRE-crYOLO is a fast and accurate fully automated particle picker for cryo-EM. Commun Biol 2, 218 (2019). https://doi.org:10.1038/s42003-019-0437-z

35 Scheres, S. H. & Chen, S. Prevention of overfitting in cryo-EM structure determination. Nat Methods 9, 853–854 (2012). https://doi.org:10.1038/nmeth.2115

36 Rosenthal, P. B. & Henderson, R. Optimal determination of particle orientation, absolute hand, and contrast loss in single-particle electron cryomicroscopy. J Mol Biol 333, 721–745 (2003). https://doi.org:10.1016/j.jmb.2003.07.013

37 Pettersen, E. F. et al. UCSF Chimera--a visualization system for exploratory research and analysis. J Comput Chem 25, 1605–1612 (2004). https://doi.org:10.1002/jcc.20084

38 Waterhouse, A. et al. SWISS-MODEL: homology modelling of protein structures and complexes. Nucleic Acids Res 46, W296–W303 (2018). https://doi.org:10.1093/nar/gky427

39 Emsley, P., Lohkamp, B., Scott, W. G. & Cowtan, K. Features and development of Coot. Acta Crystallogr D Biol Crystallogr 66, 486–501 (2010). https://doi.org:10.1107/S0907444910007493

40 Afonine, P. V. et al. Real-space refinement in PHENIX for cryo-EM and crystallography. Acta Crystallogr D Struct Biol 74, 531–544 (2018). https://doi.org:10.1107/S2059798318006551

41 Afonine, P. V. et al. New tools for the analysis and validation of cryo-EM maps and atomic models. Acta Crystallogr D Struct Biol 74, 814–840 (2018). https://doi.org:10.1107/S2059798318009324

42 Pettersen, E. F. et al. UCSF ChimeraX: Structure visualization for researchers, educators, and developers. Protein Sci 30, 70–82 (2021). https://doi.org:10.1002/pro.3943

